# Deep visual proteomics reveals distinct proximal tubular and glomerular injury programs in experimental diabetic kidney disease

**DOI:** 10.64898/2026.09.01.748754

**Authors:** Xiang Zheng, Mariavittoria d’Acierno, Lena K. Rosenbaek, Markus Rinschen, Qi Wu, Robert A. Fenton

**Affiliations:** Department of Biomedicine, Aarhus University, Aarhus, Denmark

**Keywords:** CKD, nephropathy, mass spectrometry, nephron

## Abstract

**Background:** Diabetic kidney disease (DKD) is the leading cause of chronic kidney disease (CKD). However, most proteomic studies of DKD rely on bulk kidney tissue, which cannot distinguish the contribution or response of individual nephron compartments to injury.

**Methods:** Diabetes was induced in male mice by streptozotocin (STZ) injections. After 16-weeks, the mice and vehicle-treated controls were characterized physiologically, biochemically, and histologically. A deep learning-powered Deep Visual Proteomics (DVP) pipeline, validated against manual annotation, was adapted to isolate proximal tubule (PT) and glomeruli from Megalin stained kidney sections by automated laser microdissection. Bulk kidney, PT, and glomerular proteomes were generated by data independent acquisition mass spectrometry. PT-enriched candidates were prioritized using a composite scoring approach and compared with human tubulointerstitial proteomic data from the Kidney Precision Medicine Project.

**Results:** STZ mice developed sustained hyperglycaemia and albuminuria, alongside elevated markers of tubular injury and interstitial fibrosis. Segmentation models isolated PT and glomeruli with high fidelity (Dice coefficients 0.878 and 0.914; area correlations r=0.993 and r=0.996). Compartment-resolved proteomics determined that PT and glomeruli underwent largely distinct, non-overlapping remodelling: PT exhibited loss of proteostatic, cell cycle, and structural programs with compensatory mitochondrial and lipid metabolic upregulation, whereas glomeruli showed broad loss of oxidative metabolic capacity without any compensatory metabolic program. Fourteen of the top twenty prioritized PT candidates, including LARS2 and ANXA2, changed in the same direction in human CKD tubulointerstitial proteomic data. The STZ PT proteome correlated significantly with this human dataset, while the glomerular comparison did not.

**Conclusions:** Compartment-resolved and deep learning-guided visual proteomics can uncover divergent, biologically coherent PT and glomerular injury programs in DKD that are masked in bulk tissue analysis. A PT injury signature was uncovered that is partially conserved in human CKD, identifying novel candidate mechanisms and biomarkers for future exploration.

## Introduction

Diabetic kidney disease (DKD) remains the leading cause of chronic kidney disease (CKD) and kidney failure ^1,2^. Despite substantial advances in glycaemic control, blood pressure management, and renin angiotensin system and SGLT2 inhibitor-based therapy, a large proportion of patients with diabetes still progress to advanced kidney failure ^3,4^. Population-based cohort and registry studies consistently document substantial residual risk ^5^. Current therapies therefore appear to slow, but not reverse, disease progression, highlighting major gaps in our understanding of compartment-specific kidney injury in diabetes ^4^.

The pathophysiology of DKD has traditionally been framed around the glomerulus, with mesangial expansion, glomerular basement membrane thickening, podocyte loss, and glomerulosclerosis providing the histological basis for staging systems built around albuminuria ^6^. Yet histopathological cohort studies have repeatedly shown that tubulointerstitial fibrosis correlates more closely with the rate of glomerular filtration rate (GFR) decline than glomerular lesion severity alone ^7,8^. Furthermore, single-cell and single-nucleus transcriptomic studies of human and murine diabetic kidneys have described discrete, injured proximal tubule (PT) states marked by cell cycle arrest, loss of differentiation markers, and altered metabolic gene expression ^9^. These observations have shifted the field toward viewing the PT as an active participant in disease progression rather than a passive recipient of glomerular filtrate injury. Although this shift in understanding has been based largely on dissociated single-cell datasets that are difficult to link back to intact tissue architecture, recent spatial approaches have solidified such ideas ^10–12^. However, what remains a knowledge gap is that most of these concepts are formed around RNA expression profiling, and it is not completely understood whether these compartment-specific injury programs are similarly reflected at the protein level and hence can be more readily adapted to enhance common kidney pathology grading systems.

Most kidney mass spectrometry-based proteomic studies, in contrast to transcriptomic profiling, still relies on bulk tissue, which necessarily averages the biochemical signatures of tubules, glomeruli, vasculature, and interstitium into a single composite readout. Because cellular protein abundance is shaped by translation, stability, and secretion in ways that transcript abundance alone cannot predict, this compartment blindness is a particular limitation for proteomics, where a pathway enriched in bulk tissue cannot on its own be attributed to a specific nephron segment, an infiltrating cell population, or a combination of several ^13^. However, Deep Visual Proteomics (DVP) has the capacity to overcome these limitations. DVP combines high content tissue imaging, machine learning-based image segmentation, and automated laser microdissection with sensitive data independent acquisition (DIA) mass spectrometry on routinely processed formalin fixed, paraffin embedded (FFPE) tissue ^14–16^. DVP provides a reproducible, quantitative framework for isolating any renal structure, from specific tubule segments to glomeruli, within the same kidneys used for physiological and histological phenotyping. This enables direct linkage of compartment-specific proteomic remodelling to a precisely defined disease phenotype ^17^. This strategy has begun to reshape spatial biology in oncology, where it has been used to resolve tumour and stromal proteomes at compartment or even single-cell resolution, but it has not yet been broadly applied to kidney ^15,18–21^.

In this study, we refined and applied a DVP pipeline to assess the streptozotocin (STZ)-induced model of DKD ^22^; a well-characterized model with a robust, PT and glomerular injury phenotype with measurable functional and structural correlates, providing a tractable system in which to validate compartment-resolved proteomics before extending the approach to more complex kidney disease models ^23^. We hypothesized that PT and glomeruli undergo distinct, largely non-overlapping proteomic remodelling under diabetic stress that is obscured when bulk kidney tissue is analysed alone, and that a subset of PT-specific changes identified in this model would be shared with human CKD. Our results show that compartment-resolved, deep learning-guided visual proteomics can be applied to the kidney to uncover divergent, biologically coherent PT and glomerular injury programs in DKD that are masked in bulk tissue analysis. A PT injury signature was uncovered that is partially conserved in human CKD, identifying novel candidate mechanisms and biomarkers for future exploration.

## Materials and methods

### Animal model and study design

6-week-old male C57BL/6J (Taconic) mice were subjected to five daily intraperitoneal injections of STZ (50 mg/kg body weight), freshly dissolved immediately before each injection in 0.1 M sodium citrate buffer (pH 4.5). Age and sex matched littermates received vehicle alone. Plasma glucose was measured every two weeks using a Contour Next blood glucose meter (Ascensia Diabetes Care, Parsippany, NJ, USA) and mice that became hyperglycaemic were included in the experimental cohort (n=6 per group). Mice were maintained for a further 16 weeks, with body weight recorded regularly. At the end of the study, mice were acclimatized in metabolic cages for 3 days, followed by a urine collection period of 24 h where intake of food and water, and output of urine were recorded. Subsequently, mice were anaesthetized with isoflurane before blood was sampled via the retro-orbital plexus. The left kidney was clamped, removed, cut in half transversely and rapidly frozen in liquid nitrogen. Plasma, urine and the left kidney samples were stored at -80°C prior to analysis. The right kidney was perfused with 4% paraformaldehyde (PFA) in PBS and after removal further fixed in similar buffer overnight at 4 °C before paraffin embedding. All animal procedures were approved by the Danish Animal Experiments Inspectorate (license numbers 2019-15-0201-01656, 2019-15-0201-00086 and 2024-15-0201-01672) and performed in accordance with institutional and national guidelines for the care and use of laboratory animals.

### Plasma and urine biochemistry

Plasma and urine glucose, urea, sodium, potassium, chloride, and creatinine were measured at the Clinical Pathology Laboratory of the Medical Research Council (Harwell, Oxfordshire, UK). Urine osmolality was measured using a freezing point depression osmometer (Advanced Instruments, Norwood, MA, USA; Model 3320). Urinary albumin concentration was determined using a commercially available mouse ELISA kit (Proteintech Group, Rosemont, IL, USA; Cat# KE10117).

### Semi-quantitative RT-PCR (RTqPCR)

Total RNA was extracted from half of each kidney and reverse transcribed using SuperScript™ IV CellsDirect™ cDNA Synthesis Kit (Thermo Fisher Scientific, Waltham, MA, USA). PCR was performed on a LightCycler 480 (Roche, Basel, Switzerland) and data analysed using the comparative Ct method as previously described ^24^. Primer sequences: *Kim1,* forward primer (F) 5′—CTGCTGCTACTGCTCCTTGT—3′, reverse primer (R) 5′—GCAACCACGCTTAGAGATGC—3′; *Lcn2,* F 5′—ATGTCACCTCCATCCTGGTCAG—3′, R 5′—GCCACTTGCACATTGTAGCTCTG—3′; *Cxcl10,* F 5′—CCAAGTGCTGCCGTCATTTTC—3′, R 5′—GGCTCGCAGGGATGATTTCAA—3′; *Ccl2,* F 5′—CAGGTCCCTGTCATGCTTCT—3′, R 5′—GTGGGGCGTTAACTGCATCT—3′; *Vcam1,* F 5′—GCTATGAGGATGGAAGACTCTGG—3′, R 5′—ACTTGTGCAGCCACCTGAGATC—3′; *Mrc1,* F 5′—GTTCACCTGGAGTGATGGTTCTC—3′, R 5′—AGGACATGCCAGGGTCACCTTT—3′; *18S,* F 5′—GGATCCATTGGAGGGCAAGT—3′, R 5′—ACGAGCTTTTTAACTGCAGCAA —3′;

### Histology and immunohistochemistry

Paraffin embedded kidneys were sectioned at 5 µm onto either Superfrost Plus slides (Waldemar Knittel, Braunschweig, Germany; Cat# 2510.1250 (Hounisen)) or PEN membrane slides (MicroDissect GmbH, Herborn, Germany; Cat# 11505158). Superfrost Plus mounted sections were stained with Sirius red to assess interstitial fibrosis. To delineate the PT compartment, immunohistochemistry (IHC) for the PT-specific protein Megalin was performed on PEN slides. Sections underwent deparaffinization and rehydration followed by antigen retrieval using 1% SDS for 10 min at room temperature. Endogenous peroxidase activity was blocked with 3% H₂O₂, and sections were incubated with Megalin antibody (raised against Megalin/gp330 ^25^; 1:3000) overnight at 4°C, followed by washing and incubation with HRP-conjugated rabbit anti-sheep immunoglobulins (Agilent Technologies Denmark, Glostrup, Denmark; Cat# P0163). Signal was developed using 3,3-diaminobenzidine tetrahydrochloride hydrate (Sigma-Aldrich/Merck, St. Louis, MO, USA; Cat# D6637), and sections were counterstained with Mayer’s haematoxylin solution (VWR International, Radnor, PA, USA; Q Path Cat# 10047105). Images of the whole section were collected at 40× magnification on an Olympus VS120 whole-slide scanner and stored in .vsi format to preserve acquisition metadata.

### Sirius red quantification

A pixel classifier was developed in QuPath (v0.5.0) to segment Sirius red-labelled tissue features automatically. Training was based on manual annotations from representative regions across multiple slides, with three classes defined: Sirius red positive, Sirius red negative, and background. To improve generalizability, annotations were drawn from multiple slides. The classifier was trained using QuPath’s Random Trees algorithm at 0.34 µm/pixel resolution, with Gaussian, Laplacian of Gaussian, and weighted deviation features. Training was refined iteratively by inspecting prediction maps and adding annotations in regions that were misclassified until the output was satisfactory. The final classifier was then applied to all images, and quantitative outputs, including area fraction per class, were exported for downstream analysis. Classification quality was checked by visual inspection of overlay images and, where relevant, by comparison with manual annotations or independent validation regions. Reproducibility was assessed across biological replicates.

### General statistical analysis

Data were tested for normality and variance homogeneity. Pairwise comparisons were performed using Student’s two-sided *t*-test using GraphPad Prism 10. Data are presented as mean ± SEM unless otherwise stated in legends.

### Image annotation and deep learning segmentation

To enable compartment resolved proteomics, Superfrost Plus mounted sections were additionally stained for megalin and the Na,K-ATPase (clone C464.6, Sigma-Aldrich/Merck, St. Louis, MO, USA; Cat# 05-369 ^26^; 1:1000) using a standard double-labelling protocol ^27^. Secondary antibodies were Alexa Fluor 488 donkey anti-sheep IgG (Thermo Fisher Scientific/Molecular Probes, Waltham, MA, USA; Cat# A11015; 1:1000) and Alexa Fluor 555 donkey anti-mouse IgG (Thermo Fisher Scientific/Molecular Probes, Waltham, MA, USA; Cat# A31570; 1:1000). Nuclei were counterstained with DAPI (Sigma-Aldrich/Merck, St. Louis, MO, USA; Cat# D9542). PT and glomeruli were manually annotated on whole-slide images to generate ground truth training data. Two deep learning segmentation models were trained on the arivis Cloud platform (Zeiss/arivis AG, Rostock, Germany). Model #1 segmented and classified PT and glomeruli from megalin IHC images and was used to generate the compartment masks applied in this study. Model #2 segmented tubules from megalin and Na,K-ATPase immunofluorescence images and was trained and validated. A third, pretrained cytoplasmic segmentation model integrated in BIAS software (Single-Cell Technologies, Szeged, Hungary) was combined with a machine learning classifier to resolve individual PT epithelial cells at single cell resolution ^14–16^. Both principal models were applied to full resolution images and validated on an independent tissue set prior to batch application, as described below.

### Segmentation model validation

An independent validation set of 18 kidney images, not used for model training, was manually annotated and compared separately for PT and glomeruli with the corresponding model predictions. Megalin positive regions were used as the reference standard for PT, and glomerular boundaries were annotated based on morphology and the characteristic megalin negative staining pattern. Segmentation performance was quantified as the Dice coefficient (2TP / (2TP + FP + FN)), precision (TP / (TP + FP)), and recall (TP / (TP + FN)), where TP, FP, and FN denote pixels correctly labelled as foreground (true positives, TP), pixels incorrectly labelled as foreground (false positives, FP), and foreground pixels missed by the segmentation (false negatives, FN), respectively. Agreement between manually annotated and model predicted compartment area across images was additionally assessed by Pearson correlation.

### Laser microdissection and mass spectrometry sample preparation

Model derived compartment masks were used to automate laser microdissection of PT and glomeruli from STZ and vehicle kidney sections using a Leica LMD7 with Leica LMD software (v8.5; Leica Microsystems, Wetzlar, Germany) ^15,16^. Tissue was collected in caps of 0.2 ml low protein-bind tubes and processed for mass spectrometry. To standardize sample input across specimens, mass spectrometry injection quantity was normalized to the microdissected surface area. Precursor identification plateaued above 2.3 × 10^5^ µm², which was applied as the minimum acquisition area for all samples in this study.

### Sample preparation for bulk kidney proteomics

Half of each kidney was thawed on ice and homogenized in ice cold isolation buffer (0.3 M sucrose, 25 mM imidazole, and 1 mM EDTA, pH 7.2) with protease and PhosSTOP phosphatase inhibitors (Roche Diagnostics, Mannheim, Germany; Cat# 04693124001 and 04906837001). An aliquot was reserved and archived at −80°C, and the remainder was sonicated on ice for 2 × 5 pulses at 35% tip power (QSonica, Newtown, CT, USA; Model Q125) and clarified by centrifugation at 500 *g* for 2 min. The supernatant protein concentration was determined by BCA assay (Thermo Fisher Scientific/Pierce, Waltham, MA, USA; Cat# 23225), and 50 µg of protein per sample was diluted in isolation buffer to a final volume of 180 μL. SDS was added to a final concentration of 1%. Samples were sonicated again (2 × 5 pulses, 35% tip power) and clarified by centrifugation at 16,000 *g* for 10 min. Cleared lysates were processed by filter aided sample preparation (FASP) and desalted prior to mass spectrometry analysis as described ^28^.

### Sample preparation for compartment resolved (PT and glomerular) proteomics

Microdissected PT and glomerular samples were processed using a low input, on bead digestion workflow. Tubes were briefly centrifuged at 16,000 *g* and 5 μL of 100 mM triethylammonium bicarbonate (TEAB) buffer was added to the lid before a second centrifugation step for 1 min. After adding an extra 150 μL of 100 mM TEAB, samples were heated at 95°C for 5 min, immediately chilled on dry ice for 5 min, then centrifuged and sonicated on ice using short pulse cycles (10 cycles, 5 s on/5 s off) to promote tissue disruption. Proteins were reduced with 10 mM DTT in 8 M urea, 100 mM TEAB buffer for 1 h at 37°C, then alkylated in the dark with 100 mM iodoacetamide in TEAB for 30 min at room temperature. Samples were proteolytically digested sequentially by adding 10 μL of 100 ng/nL Lys-C at 37 °C for 2 hours, followed by adding 10 μL of 200 ng/nL of trypsin overnight at 37 °C. Digestion was quenched with adding 5 μL of 10% trifluoroacetic acid (TFA) and peptides were desalted using C18 StageTips (Thermo Fisher Scientific/Savant, Waltham, MA, USA) following standard procedures. Eluates were dried by SpeedVac at 60°C prior to mass spectrometry analysis.

### Liquid chromatography mass spectrometry (LC-MS/MS)

Peptides were separated on a Thermo Scientific Vanquish LC system equipped with a Micropep column (Thermo Fisher Scientific, Waltham, MA, USA) using a 30-minute gradient with an active separation window of 22 min, ramping from 5 to 22% solvent B (80% acetonitrile with 0.1% formic acid) over 17 min and then from 22 to 40% over the following 5 min. Subsequent MS/MS analysis was conducted on an Orbitrap Ascend mass spectrometer (Thermo Fisher Scientific, Waltham, MA, USA) equipped with a field asymmetric ion mobility spectrometry (FAIMS) Pro Duo interface. Data were acquired in data independent acquisition (DIA) mode with FAIMS compensation voltage fixed at −45 for both MS1 and MS2. MS1 scans spanned an m/z range of 375 to 1010 at an Orbitrap resolution of 240,000, with a maximum injection time of 507 ms and a normalized AGC target of 250%. MS2 scans used a 68 Da isolation window across an m/z range of 400 to 1000, at matching Orbitrap resolution and injection time settings, a normalized AGC target of 1000%, and normalized collision energy of 32%, with nine scan events and a loop count of three per cycle.

### Mass spectrometry data processing

Raw files were processed in Spectronaut (v18; Biognosys AG, Schlieren, Switzerland) using direct DIA+ search against a mouse UniProt reference proteome (proteome ID UP000000589, downloaded January 6, 2025), with peptide and protein false discovery rates set at 1%. Quantification was performed at the MS1 level throughout. For searches default settings were used except removing carbamidomethylation of cysteine from the fixed modification panel. Identified proteins and their corresponding quantification matrices were exported as .CSV files for downstream statistical analysis.

### Proteomics data processing, filtering, and imputation

Label free quantification intensities were processed in Perseus (v2.0.10.0; Max Planck Institute of Biochemistry, Martinsried, Germany), log2 transformed, and filtered ^29^. For differential abundance testing, the intensity matrix was filtered to retain proteins with non-missing values in at least 70% of samples in at least one of the two experimental groups (STZ or vehicle). For multivariate analyses requiring a complete matrix, including principal component analysis (PCA) and clustering, missing values were imputed using Perseus default Gaussian distribution-based imputation (width 0.3, downshift 1.8 standard deviations), an approach commonly used to approximate low abundance, left censored signal in label free proteomics ^15^. Imputation was evaluated by comparing observed and imputed intensity distributions, assessing the stability of differential expression results with and without imputation, and confirming that the identity of the top differentially abundant proteins was robust to alternative imputation parameters.

### Differential abundance and statistical analysis of proteomic data

Two sample t tests, with Benjamini-Hochberg correction for multiple testing, were performed on the filtered, unimputed, log2 transformed intensity matrix to compare STZ and vehicle groups within each compartment. Log2 fold change was calculated as the difference between mean log2 intensity in the STZ and vehicle groups. No protein retained significance at 5% FDR in any comparison; results were therefore treated as exploratory, and differentially abundant proteins were defined for downstream analysis and pathway enrichment using an unadjusted P value below 0.05 combined with an absolute fold change of at least 1.5, prioritizing proteins with large, consistent effect sizes across compartments for further interpretation.

### Pathway enrichment analysis

Pathway enrichment was performed in Cytoscape (version 3.10.2) using the ClueGO plugin (version 2.5.10). Significantly upregulated and downregulated proteins from each comparison were analysed separately as gene symbol lists against Gene Ontology Biological Process terms and curated pathway databases, including KEGG and Reactome, using a right sided hypergeometric test with Benjamini-Hochberg correction (FDR < 0.05) ^15^. Functionally related terms and pathways were grouped based on kappa statistics, and representative, non-redundant terms from each cluster were used to summarize enriched biological processes.

### PT candidate prioritization

Candidate PT markers were prioritized by integrating differential proteomics results from the PT, glomerular, and bulk kidney comparisons (STZ versus vehicle) with pathway annotations. Protein identifiers were standardized to gene symbols, and duplicate entries within each dataset were summarized by mean log2 fold change and maximum significance score. Effect strength was defined as 0.75 × |PT log2FC| + 0.25 × (−log10 P_PT), and PT specificity was defined as |PT log2FC| − (|glomerular log2FC| + |bulk log2FC|)/2; both terms were standardized by z score transformation across candidates. Pathway relevance was calculated as the number of enriched pathways containing each gene, capped at three and normalized to a 0 to 1 scale. Biomarker feasibility was assigned manually on a 0 to 3 ordinal scale, with higher scores given to secreted, extracellular, membrane associated, or shed proteins based on expected detectability and translational plausibility. The composite prioritization score was calculated as 0.35 × EffectScore + 0.30 × SpecificityScore + 0.20 × PathwayScore + 0.15 × FeasibilityScore. Weights were assigned a priori to reflect the relative importance of each criterion for identifying clinically translatable proximal tubule injury markers: effect strength and compartment specificity were weighted most heavily, because a candidate biomarker must first show a robust, PT-selective response to injury, whereas pathway relevance and biomarker feasibility were weighted lower as supportive rather than primary criteria, contextualizing rather than establishing candidate relevance. To assess the robustness of this ranking, we confirmed that the top-ranked candidates remained largely stable across a range of alternative weightings (**Supplementary Table 1**). Candidates were ranked in descending order of this score, with regulation direction retained separately based on the sign of the PT log2 fold change.

### Human kidney proteomic data and cross species comparison

Human tubulointerstitial and glomerular proteomic data were obtained from the KPMP Atlas Explorer v1.0 regional proteomics dataset (DOI: 10.48698/6vrn-ze53 and 10.48698/te4e-e195), generated by the KPMP Kidney Tissue Atlas ^30,31^. This resource comprises aggregated, sub-segmentally laser-microdissected proteomic profiles from human kidney biopsies and nephrectomy tissue spanning kidney injury (AKI, n=13) or CKD (n=13) diagnosis. All data were accessed under KPMP’s open access data use terms (Accessed 19 December 2023. https://www.kpmp.org). Log2 fold changes for all genes in the mouse and human datasets were independently z score normalized. Mouse genes were mapped to human orthologs using the UniProt ID Mapping tool, and only genes shared between species were retained for comparison. Spearman correlation coefficients were calculated between the normalized log2 fold changes of shared genes, separately for the PT and glomerular comparisons, to quantify similarity between the STZ model and human tubulointerstitial CKD proteome. Spearman correlation was chosen for its robustness to outliers and its suitability for monotonic relationships that are not linear.

## Results

### Mouse STZ-induced diabetes recapitulates key DKD hallmarks

STZ-treated mice developed sustained hyperglycaemia within two weeks of injection, with plasma glucose remaining approximately two-fold higher than vehicle-treated mice (∼20 mmol/L) for the study period (**Figure 1A**). Consistent with overt diabetes, after 16 weeks the STZ-treated mice had significantly lower body weight despite unchanged food intake, and significantly increased water intake and urine output (**Figure 1B**; **Supplementary Figure 1A**). STZ-treated mice showed markedly higher urinary glucose excretion, significantly increased urinary sodium and potassium excretion, and the albumin to creatinine ratio (UACR) was raised (**Figure 1B**). However, creatine clearance was not different between groups (**Supplementary Figure 1A)**.

**Figure 1.**
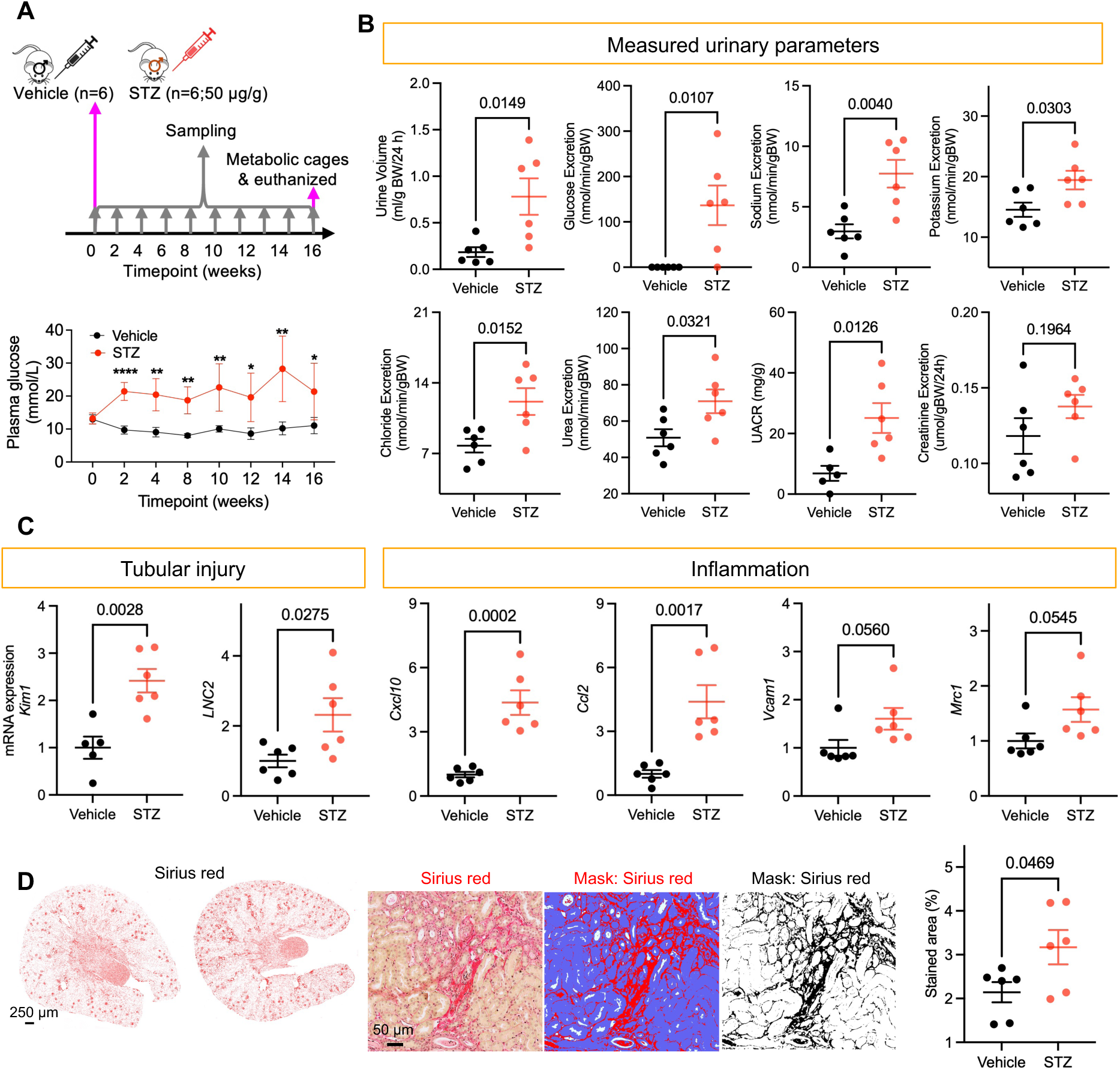
Mouse STZ-induced diabetes recapitulates key hallmarks of diabetic kidney disease. (A) Study design and plasma glucose concentrations throughout the 16-week study in vehicle and STZ-treated mice (n=6 per group). Statistical analyses were performed using a two-way ANOVA followed by Tukey multiple comparison testing. P values between conditions are denoted as follows: *P<0.05, **P<0.01, ***P<0.001, ****P<0.0001. (B) Urine volume, urinary excretion of glucose, sodium, potassium, chloride, urea, and creatinine, and urinary albumin to creatinine ratio (UACR) in vehicle and STZ-treated mice. (C) Relative kidney mRNA transcript levels of *Kim1*, *Lcn2*, *Cxcl10*, *Ccl2*, *Vcam1*, and *Mrc1* determined by RTqPCR. (D) Representative images of Sirius Red stained kidney sections showing interstitial fibrosis in vehicle and STZ-treated mice. Graphs in (B) and (C) show mean ± SEM and individual data points represent values from an individual mouse. Statistical significance was determined using an unpaired two-tailed Student’s *t test*.

Relative to vehicle controls, RTqPCR gene expression analysis showed higher mRNA levels of *Kim1* (encoding Kidney Injury Molecule-1), *Lcn2* (encoding Lipocalin-2/NGAL), *Cxcl10 (*encoding the C-X-C motif chemokine ligand 10 protein), *Ccl2* (encoding the Monocyte Chemoattractant Protein-1), *Vcam1* (encoding Vascular Cell Adhesion Molecule-1) and *Mrc1* (encoding the Macrophage Mannose Receptor 1) in STZ-kidneys (**Figure 1C**). These findings indicate that the STZ-model has classic molecular signatures of injured and inflamed kidney tissue with PT injury and active recruitment of immune cells (monocytes/macrophages/T-cells/NK-cells)^32,33^.

The higher mRNA levels of *Mrc1* in kidneys of STZ-treated mice suggests macrophage infiltration and transition towards fibrosis development ^34^. To confirm this, collagen levels were assessed by Sirius Red histological staining and quantification. Kidneys from STZ-treated mice had Sirius Red staining over a significantly larger area of the kidney compared with vehicle controls (**Figure 1D**), confirming development of fibrosis.

Taken together, the data support that the utilized STZ model has several hallmarks of early DKD, characterised by sustained hyperglycaemia, systemic manifestations of diabetes, altered tubular solute handling, selective albuminuria with activation of PT injury and inflammatory pathways, and early interstitial fibrosis.

### DVP pipeline enables accurate and reproducible isolation of PT and glomeruli

To isolate PT and glomeruli from the same kidney sections for compartment-resolved proteomic profiling, we applied a DVP pipeline (**Figure 2A**). Megalin immunohistochemistry (IHC) was used to identify PT, and glomeruli were identified by their characteristic structure and negative staining (**Figure 2B, C**; **Supplementary Figure 1B**). A deep learning segmentation model was trained on manually annotated PT and glomeruli to generate compartment masks that directed automated laser microdissection (Model #1; **Figure 2A, C**). To make the pipeline adaptable to alternative staining strategies, a second segmentation model based on Megalin and Na,K-ATPase immunofluorescence labelling was benchmarked, trained and validated as a staining agnostic alternative (Model #2; **Figure 2C**).

**Figure 2.**
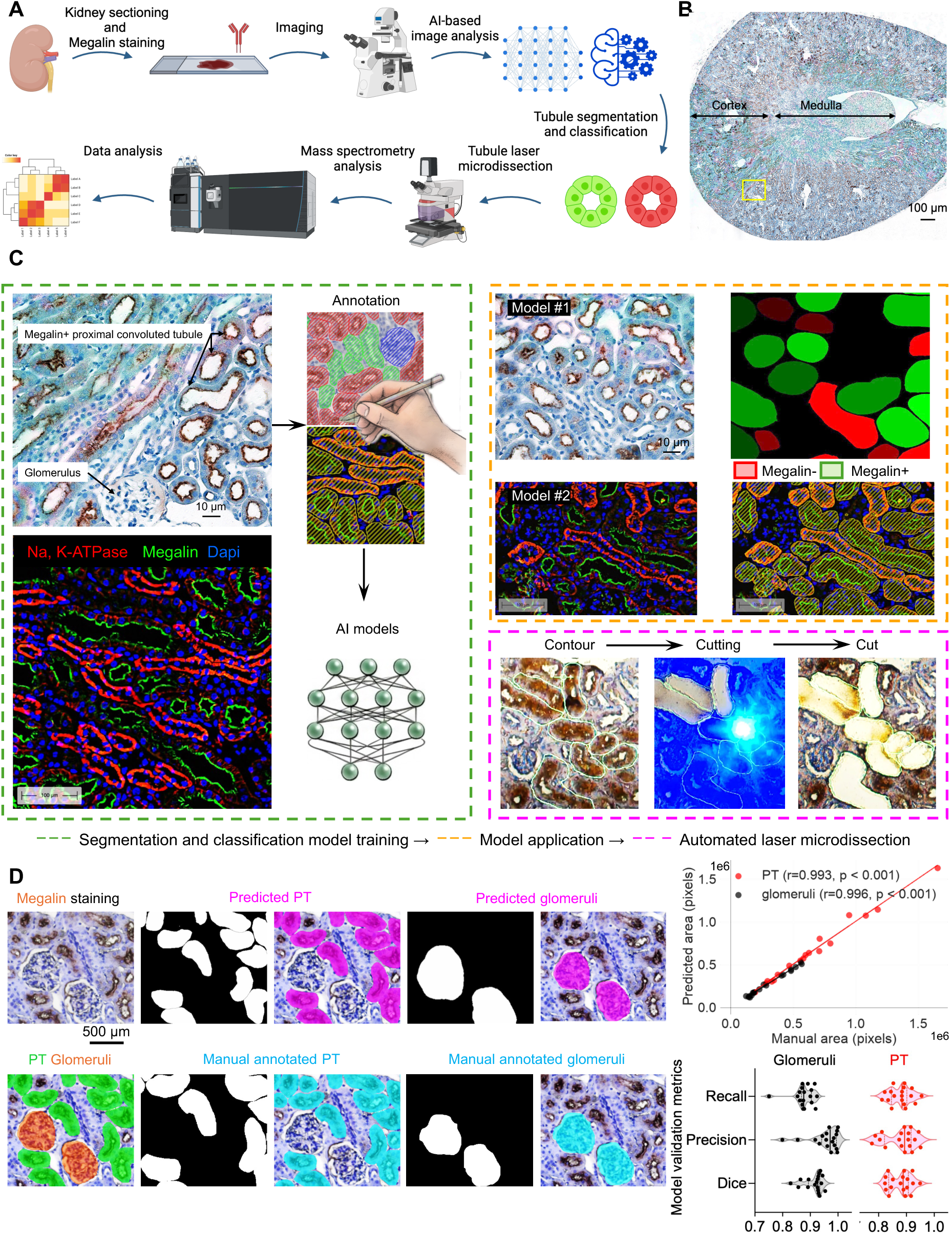
Deep visual proteomics pipeline enables accurate isolation of PT and glomeruli. (A) Schematic of the Deep Visual Proteomics (DVP) workflow, in which immunohistochemistry guided image segmentation directs automated laser microdissection of PT and glomeruli from kidneys for downstream proteomic profiling. (B) Representative whole kidney Megalin immunohistochemistry image; boxed region corresponds to region in panel C. (C) Higher magnification of the boxed region showing Megalin positive PT and Megalin negative glomeruli. Deep learning models trained on manually annotated tubules and glomeruli generated compartment masks (dotted orange box) for automated laser microdissection (dotted pink box). (D) Segmentation model performance (predicted, magenta) relative to manual annotation (cyan). Dice coefficient, precision, and recall quantify pixel level spatial overlap between predicted and manually annotated compartments; Pearson correlation (r) quantifies agreement between manually annotated and predicted compartment area across images (See **Methods**).

Segmentation performance was benchmarked against manual annotation using independent images excluded from model training. The PT model achieved a mean Dice coefficient of 0.878 ± 0.039, precision of 0.879 ± 0.039, and recall of 0.879 ± 0.039, with the predicted and manually annotated PT area strongly correlating (r=0.993, p<0.001). The glomerular model performed even better on overlap metrics, with a Dice coefficient of 0.914 ± 0.034, precision of 0.956 ± 0.053, recall of 0.878 ± 0.038, and an area correlation of r=0.996 (P<0.001), indicating high specificity with a modest reduction in sensitivity (**Figure 2D**). A complementary single-cell segmentation and classification model further resolved individual PT epithelial cells for future single cell applications (Model #3; **Supplementary Figure 2A**). To standardize mass spectrometry input across samples, we compared the microdissected surface area with precursor ion quantity and identified a plateau (∼2.3 × 10^5^ µm²) where greater sample input did not increase the number of precursors identified (**Supplementary Figure 2B**). This acquisition area was subsequently used for all samples in the STZ analysis study. Together, these benchmarks demonstrate that the DVP pipeline isolates PT and glomeruli with high spatial fidelity and reproducibility, providing a validated foundation for compartment-resolved proteomic profiling.

### Compartment-resolved proteomics reveals divergent PT and glomerular alterations in DKD

Bulk kidney, plus microdissected PT and glomeruli were each profiled by DIA mass spectrometry, yielding approximately 5,000, 2,000, and 2,500 protein groups respectively. Roughly 3,000 proteins were identified across the combined PT and glomerular datasets (**Figure 3A**; **Supplemental Data 1**).

**Figure 3.**
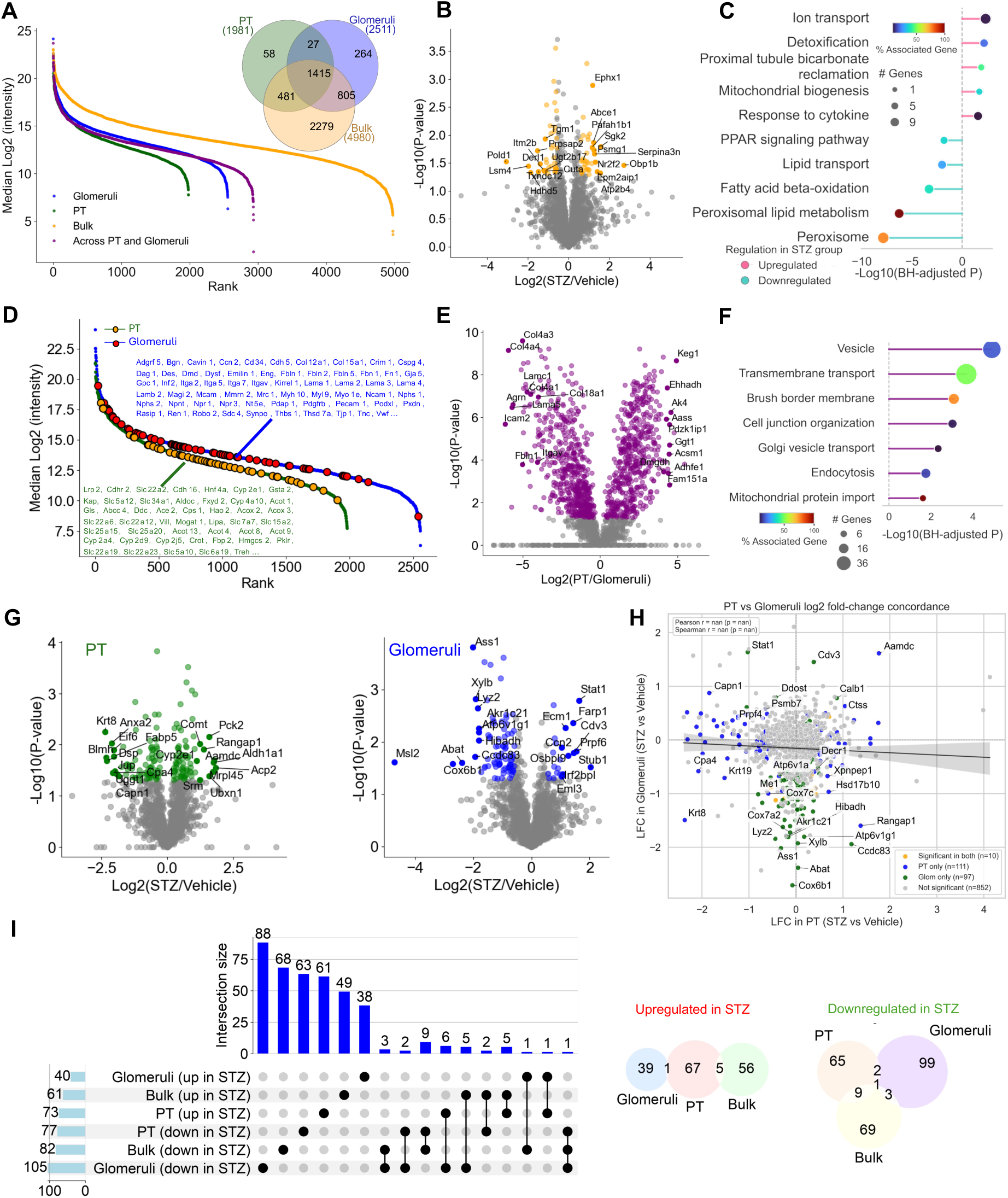
Compartment resolved proteomics reveals divergent PT and glomerular alterations in DKD. (A) Rank abundance plots of protein groups identified in bulk kidney, glomeruli, and PT, and protein numbers identified across the combined PT and glomerular datasets. (B) Volcano plot illustrating proteomic comparison of bulk kidney from STZ-treated versus vehicle mice. The top 20 differentially regulated proteins are highlighted. (C) Gene ontology and pathway enrichment for significantly changed proteins in bulk kidney. (D) Rank abundance comparison of PT and glomerular proteomes in kidneys from vehicle-treated mice show canonical compartment markers for PT, including Lrp2, Slc34a1, Slc22a6, Cyp2e1 and glomeruli, including Nphs1, Nphs2, Podxl, Pecam1. (E) Volcano plot illustrating proteomic comparison of PT and glomeruli in kidneys from vehicle-treated mice. The top 20 differentially regulated proteins are highlighted. (F) Pathways enriched in the PT relative to glomeruli. (G) Volcano plot illustrating proteins significantly altered in PT and glomeruli from STZ-treated versus vehicle mice. (H) Overlap of proteins significantly altered in both PT and glomeruli. (I) UpSet and venn diagram of significantly changed proteins across bulk kidney, PT, and glomeruli.

In bulk kidney tissue from STZ-treated mice, 143 proteins were significantly changed relative to vehicle (61 increased in abundance, 82 down; fold changes 1.5 to 32). Pathway analysis revealed upregulation of cytokine-responsive and stress-adaptive programs, including response to cytokine and detoxification pathways, together with mitochondrial biogenesis and ion transport modules. Notably, proteins associated to PT bicarbonate reclamation were also increased, consistent with altered tubular transport in DKD. In contrast, pathways related to peroxisomal metabolism, PPAR signalling, fatty acid beta-oxidation, lipid transport, and steroid/lipid metabolic homeostasis were reduced, indicating a broad suppression of lipid-catabolic programs in the kidney during DKD (**Figure 3B, C**). These changes suggest that STZ-treatment induces an overall shift in the kidney towards inflammatory and stress-response signalling with concurrent failure of peroxisome-linked lipid oxidation and metabolic homeostasis ^35–37^.

Not unexpectedly, analysis of PT and glomeruli from vehicle-treated mice showed clear proteomic separation, with 1,056 proteins differing between compartments (fold changes 1.5 to 64) and clear enrichment of canonical compartment markers, including *Lrp2*, *Slc34a1*, *Slc22a6*, and *Cyp2e1* in PT and *Nphs1*, *Nphs2*, *Podxl*, and *Pecam1* in glomeruli (gene symbols of corresponding proteins shown, **Figure 3D-E**). The PT enriched fraction was dominated by pathways governing mitochondrial protein import, Golgi vesicle transport, brush border membrane organization, endocytosis, vesicle trafficking, and transmembrane transport, consistent with the apical reabsorptive specialization of PT epithelium and distinct from the structural and filtration related programs of glomeruli (**Figure 3F**). This baseline separation confirmed that the microdissection strategy preserved compartment-specific biology and provided the reference frame for interpreting diabetes-induced changes ^17,38–40^.

When PT and glomeruli were analysed separately in response to STZ-treatment, each compartment showed a distinct and largely non-overlapping response (**Figure 3G-I**). The compartment specific nature of these responses was underscored by their limited overlap: only 10 proteins were altered similarly in both PT and glomeruli, versus 111 changes unique to PT and 97 unique to glomeruli (**Figure 3H**). Across bulk kidney, PT and glomeruli, no protein was commonly increased following STZ-treatment, while only Gatd3, a mitochondrial deglycase, was commonly downregulated (**Figure 3I**). GATD3 protects mitochondria from glycation-related damage, so a reduction in GATD3 activity may promote the mitochondrial dysfunction associated with DKD and is a novel mechanistic candidate for further investigation.

In PT, 150 proteins were significantly different between STZ-treated and vehicle controls (73 higher, 77 lower; fold changes 1.5 to 8). Pathway analysis indicated that in PT, STZ-induced diabetes is associated with coordinated downregulation of multiple stress-responsive and signalling modules together with structural and proliferative programs. Proteins belonging to the proteasome core complex and the cell cycle were lower in STZ-treated PT, indicating suppression of ubiquitin–proteasome-mediated protein homeostasis and mitotic/DNA-replication machinery ^41^. Inflammatory and cytokine signalling pathways, including TNF-alpha–NF-κB signalling and cellular responses to interleukin-4, were reduced, as were signalling by WNT, Hedgehog, and MAPK family cascades, pointing to broad attenuation of immune, morphogen, and stress-activated signalling ^42^. Pathways related to cellular responses to hypoxia, and supramolecular fiber organization were also reduced, consistent with impaired hypoxic adaptation and remodelling of cytoskeletal and epithelial junctional structures in diabetic PT. In contrast, pathways associated with mitochondrial and intermediary metabolism, including mitochondrial matrix, carboxylic acid metabolic process, glyoxylate and dicarboxylate metabolism, and fatty acid ω-oxidation, were increased in PT from STZ-treated mice (**Figure 3G**; **Figure 4A**).

**Figure 4.**
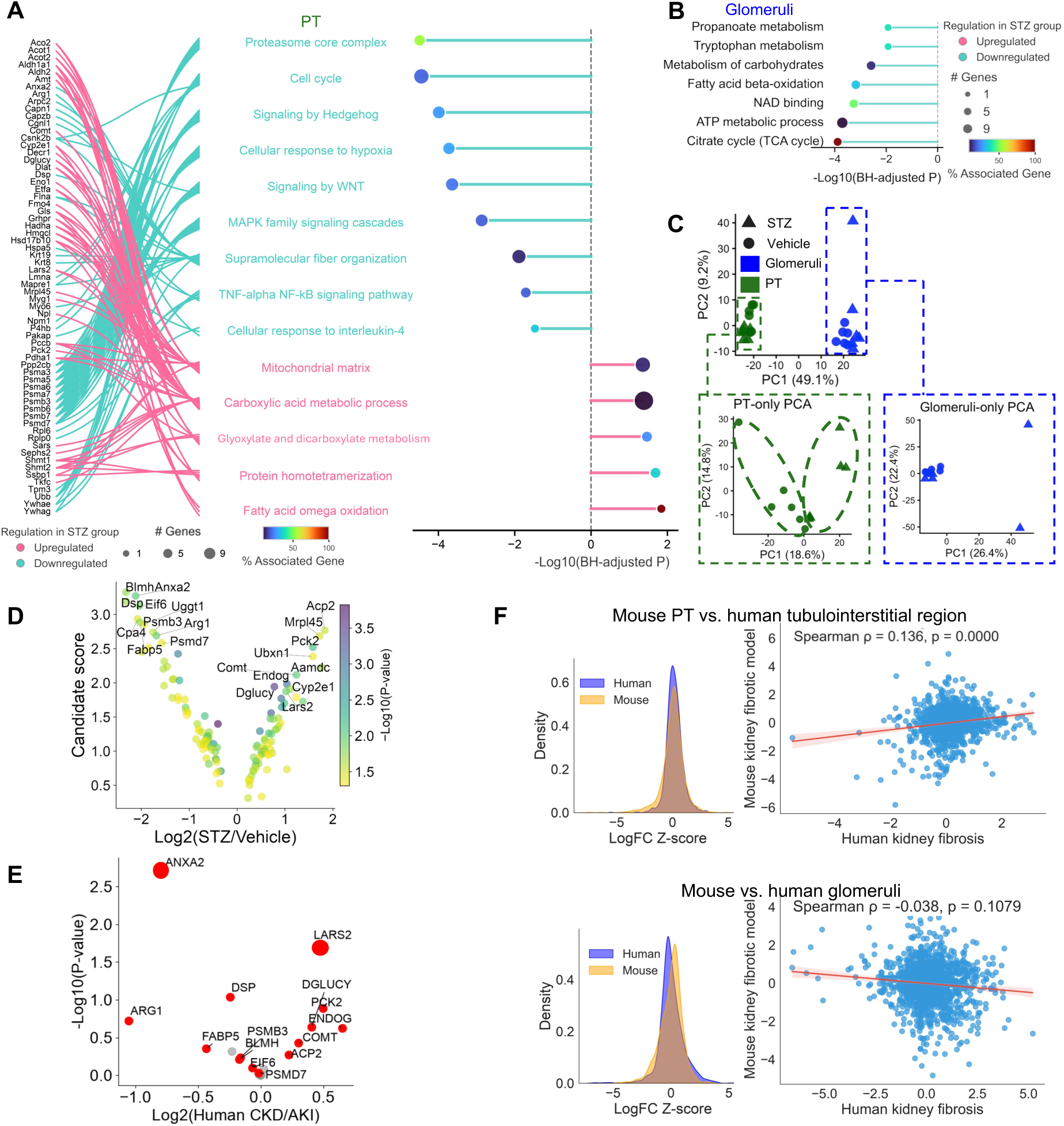
A PT enriched injury signature correlates with human chronic kidney disease. (A-B) Pathway enrichment in PT and glomeruli. In (A), sankey diagram (left) connects genes to significantly enriched pathways (right), with ribbon colors indicating direction of regulation. Differential abundance in all panels was defined by two sample *t test* (P<0.05) and an absolute fold change ≥1.5. Pathway enrichment was conducted in ClueGO, applying Benjamini–Hochberg correction and retaining terms with FDR < 0.05. (C) PCA of PT and glomeruli samples (STZ vs. vehicle). (D) PT enriched candidates ranked by a composite prioritization score integrating PT fold change, PT specificity relative to glomeruli and bulk kidney, pathway relevance, and biomarker feasibility (see Methods). Top 10 ranked increased and decreased candidates are shown. (E) Direction of change for the top 20 prioritized candidates in human chronic kidney disease data from the Kidney Precision Medicine Project. Red points indicate candidates with concordant direction of change between the STZ mouse and human datasets (14 of 20 candidates). (F) Spearman correlation of z scored log2 fold changes between the STZ proteome and the human KPMP tubulointerstitial regional proteomic dataset, shown separately for PT and glomerular comparisons.

In glomeruli, 145 proteins were significantly altered in STZ-treated mice (40 higher, 105 lower; fold changes 1.5 to 23). The proteins increased in abundance were not uniformly associated with any specific pathways, whereas the reduced proteins were associated with NAD-linked oxidative metabolism, fatty acid beta oxidation, the TCA cycle, carbohydrate metabolism, and ATP metabolic processes (**Figure 3G**; **Figure 4B**). This pattern points to a glomerular response dominated by loss of mitochondrial oxidative capacity without a compensatory metabolic program^35,43,44^.

### A PT-enriched injury signature correlates with human CKD

To define the global architecture of the STZ-induced proteomic response, we performed principal component analysis (PCA) across all PT and glomerular samples. Compartment identity accounted for the largest proportion of proteomic variance (PC1, 49.1%), clearly separating PT and glomerular proteomes, while treatment status contributed as a secondary axis of variation (PC2, 9.2%). These findings highlight the strong molecular distinction between the different compartments and indicate that diabetes-associated remodelling occurs within a pre-existing compartment-specific proteomic landscape. Focusing specifically on PT samples, PCA revealed clear segregation between STZ and vehicle groups (PC1, 18.6%; PC2, 14.8%), demonstrating a coordinated proteomic response across the PT compartment following STZ-treatment. In contrast, glomerular samples showed no apparent separation between STZ and vehicle groups (PC1, 26.4%; PC2, 22.4%), suggesting a more limited or heterogeneous glomerular proteomic response at this disease stage (**Figure 4C**).

Given the pronounced PT-centred changes observed in response to STZ-induced diabetes, we next investigated whether these molecular alterations reflected disease-relevant processes conserved in human kidney pathology. PT-enriched candidates were ranked using a composite score integrating PT fold change, PT specificity relative to glomeruli and bulk kidney, pathway relevance, and biomarker feasibility (see ***Methods***). Across 220 alternative weighting scenarios (individual weight perturbations of ± 0.05–0.10 and 200 random draws from the weight simplex), the top 20 ranked candidates showed a median overlap of 18/20 with the base ranking, a median Jaccard similarity of 0.818, and a median Spearman rank correlation of 0.989 (**Supplementary Table 1**), indicating that the composite ranking is robust to reasonable variation in the assigned weights. This approach highlighted a set of top ranked upregulated candidates, including *Acp2*, *Mrpl45*, *Pck2*, *Ubxn1*, *Aamdc*, *Comt*, *Endog*, *Dglucy*, *Cyp2e1*, and *Lars2*, and downregulated candidates, including *Blmh*, *Anxa2*, *Dsp*, *Eif6*, *Cpa4*, *Fabp5*, *Uggt1*, *Psmb3*, *Arg1*, and *Psmd7*, spanning mitochondrial metabolism, proteostasis, and epithelial structural remodelling (**Figure 4D**).

We then compared the direction of change for these top candidates with human proteomic data from the KPMP (see **Methods**), which profiles the tubulointerstitial compartment in patients with CKD of mixed and unspecified etiology, including but not limited to DKD. Fourteen of the top 20 PT candidates changed in the same direction in the human dataset, with LARS2 among upregulated proteins and ANXA2 among downregulated proteins. Five candidates showed discordant changes, and one was undetected in the human dataset (**Figure 4E**). Extending this comparison across the shared mouse to human orthologous proteome indicated that the PT proteome from STZ mice positively correlated with the human CKD tubulointerstitial proteome, whereas the corresponding glomerular comparison did not reach significance (**Figure 4F**). Together, these results indicate that the STZ model captures a PT injury program that is, at least in part, conserved in human CKD more broadly, supporting its use as a preclinical platform for investigating tubule-specific mechanisms of PT injury.

## Discussion

This study combined a well-characterized model of insulin-deficient diabetes with a deep learning-powered DVP pipeline to resolve compartment proteomic consequences of diabetic kidney injury. Three major findings emerge. First, the segmentation and microdissection pipeline isolated PT and glomeruli with a high degree of spatial fidelity. Dice coefficients exceeding 0.87 for both compartments compare favourably with deep learning-based segmentation reported in other tissues and were achieved using routinely processed FFPE sections compatible with archival and clinical workflows ^18,45^. Second, PT and glomeruli underwent largely distinct, non-overlapping proteomic remodelling under diabetic stress. PT showed a mixed injury and metabolic adaptation program, whereas glomeruli showed a more uniform loss of oxidative metabolic capacity without compensatory upregulation. Third, a subset of PT enriched proteins prioritized from this analysis showed concordant direction of change in human tubulointerstitial proteomic data from CKD patients, suggesting that at least part of the murine tubular injury program is conserved across species.

The PT proteome supports an active role for tubular epithelium in DKD rather than a secondary response to glomerular injury ^46,47^. STZ treatment reduced proteins associated with proteasomal function, cell-cycle control, cytoskeletal organisation, and cell–cell contacts while increasing proteins involved in mitochondrial and lipid metabolism. This combination is consistent with an injured epithelial state under sustained energetic pressure. Tubulointerstitial fibrosis correlates more closely with declining kidney function than the severity of glomerular lesions in histopathological cohorts ^7,8^, and single-cell and single-nucleus studies have identified injured or failed-repair PT states characterized by cell-cycle arrest, loss of differentiation, and metabolic reprogramming ^48,49^. The present data extend these observations to the proteome and show that structural injury coexists with increased abundance of mitochondrial and intermediary-metabolism proteins within the same anatomical compartment.

This metabolic enrichment requires cautious interpretation. Proteomic pathway analysis measures relative protein abundance, not mitochondrial flux or respiratory competence. One plausible explanation is an early compensatory response to increased energy demand ^50^. Hyperglycaemia, increased filtered glucose load, and hyperfiltration raise the PT reabsorptive burden and are expected to increase ATP demand. The observed proteomic profile may therefore represent an “overworked mitochondria” phase in which tubular cells expand metabolic capacity before progressing to the mitochondrial dysfunction associated with established DKD ^51^. Such compensation may initially sustain solute transport and epithelial repair but become maladaptive under persistent stress, promoting oxidative injury, lipid accumulation, and profibrotic signalling ^52–55^. A single 16-week time point cannot distinguish transient adaptation from the onset of metabolic failure. Nonetheless, the coexistence of structural injury and metabolic activation identifies PT metabolism as a candidate driver of disease progression, not merely a reflection of glomerular dysfunction ^52–55^.

The consistent reduction in GATD3 adds a potential mechanistic link between hyperglycaemia and mitochondrial injury. GATD3 was the only protein downregulated across bulk kidney, PT, and glomerular analyses following STZ treatment. As a mitochondrial deglycase implicated in protection against glycation-related damage and advanced glycation end-product accumulation, reduced GATD3 is consistent with weakened mitochondrial glycation defence across renal compartments ^56^. Loss of this protective activity could lower the threshold at which a compensatory, high-output metabolic state becomes dysfunctional ^57^. This sequence remains hypothetical. Direct measurements of mitochondrial respiration, ATP production, redox balance, reactive oxygen species, and protein glycation will be required to determine whether GATD3 loss precedes or contributes to metabolic failure in DKD.

Glomeruli followed a different trajectory to PT. STZ treatment reduced proteins involved in NAD-dependent oxidation, fatty-acid β-oxidation, the tricarboxylic acid cycle, and ATP metabolism, without a corresponding upregulated metabolic programme. This profile is compatible with impaired glomerular bioenergetics rather than structural collapse alone ^58^. Reduced oxidative capacity in podocytes has been linked to cytoskeletal instability and foot-process effacement before overt glomerulosclerosis ^44,59,60^. The glomerular proteome may therefore capture metabolic dysfunction preceding the structural damage that dominates later disease. Cell-type attribution remains limited, however, because the microdissected glomerular compartment contains podocytes, endothelial cells, mesangial cells, and associated matrix.

Compartment resolution was essential to uncover these divergent responses. Bulk kidney profiling yielded inflammatory, transport, and lipid-metabolic signatures that could not be assigned confidently to a nephron segment or distinguished from vascular, interstitial, or infiltrating-cell contributions. DVP localized bicarbonate reclamation pathway enrichment to PT and revealed that relatively few regulated proteins were shared among bulk kidney, PT, and glomerular analyses. This limited overlap is biologically informative: diabetic kidney injury is strongly compartmentalized, not a uniform tissue-wide response. This finding aligns with single-cell and spatial transcriptomic studies showing that renal injury programmes are often restricted to particular cell types or anatomical niches. Proteomic resolution adds another layer, because post-transcriptional regulation, protein turnover, and pathway activity may not follow RNA abundance.

The developed workflow also has practical methodological value. Immunohistochemistry-guided deep learning segmentation, automated microdissection, and deep proteomic analysis were implemented on standard FFPE material ^14–16^. This combination permits retrospective analysis of histologically defined compartments in archived specimens without requiring fresh tissue or tissue dissociation. The validated segmentation models, together with the stain-agnostic alternative developed here, should facilitate extension to additional nephron segments, renal cell populations, and disease models. Surface area-based sampling further standardizes input across structures of markedly different size and cellularity.

Our data comparison with human KPMP tubulointerstitial proteomic data provided a partial translational anchor. Fourteen of the 20 prioritized PT candidates changed in the same direction in human CKD. This concordance supports a composite prioritization strategy integrating effect size, compartment specificity, pathway relevance, and biomarker feasibility ^12,17^. It does not establish DKD specificity. The available human cohort comprised CKD of mixed or incompletely specified aetiology without sufficiently detailed diabetes annotation, and the tubulointerstitial samples contained both tubular epithelium and surrounding interstitium. The conserved signature may therefore represent a broader tubular injury programme involving fibrosis, cell-cycle arrest, and metabolic reprogramming rather than a diabetes-specific response. Diabetes-annotated cohorts with detailed renal, metabolic, and glycaemic phenotyping will be needed to resolve this distinction. Notably, the corresponding glomerular analysis showed no significant correlation with human data. This compartmental difference argues against uncritical extrapolation of either the STZ model or the analytical framework to glomerular-centred disease and indicates that the present model is best suited to investigating tubular injury and tubulointerstitial fibrosis.

This study has several strengths: compartment-matched analysis of bulk kidney, PT, and glomeruli within the same animals; rigorously validated segmentation; standardized microdissection; and direct cross-species comparison. However, important limitations remain. Firstly, the STZ model captures insulin-deficient hyperglycaemia but not the obesity, insulin resistance, dyslipidaemia, and cardiovascular comorbidity typical of most human DKD. Furthermore, our sample size was modest, and as individual protein remained significant after correction for multiple testing, differentially abundant proteins should be treated as hypothesis-generating until orthogonally validated. Importantly, the single 16-week time point cannot separate early drivers from adaptive or downstream responses. From a technical viewpoint, the microdissected compartments also remain multicellular, with glomerular samples containing several resident cell types and PT samples potentially containing some adjacent interstitial cells. Finally, proteomic abundance does not establish pathway activity or causal function.

In conclusion, deep learning-guided, compartment-resolved DVP revealed distinct proteomic responses in PT and glomeruli that were obscured in bulk kidney tissue. PT combined structural injury with increased abundance of mitochondrial and lipid-metabolic proteins, whereas glomeruli showed broad suppression of oxidative metabolism. Part of the tubular signature was conserved in human CKD, although its specificity to diabetic disease remains to be established. The resulting workflow provides a practical and extensible framework for resolving anatomically restricted protein programmes in standard FFPE kidney tissue and should be applicable to additional nephron compartments, experimental models, and archival human biopsies. Future work should prioritize orthogonal validation of the highest-ranking proteins, particularly GATD3 and the PT metabolic candidates, using targeted proteomics, immunostaining, and functional assays. Earlier and later time points will be required to define the proposed transition from compensatory metabolic activation to mitochondrial dysfunction.

## Data Availability

The mass spectrometry proteomics data have been deposited to the ProteomeXchange Consortium via the PRIDE partner repository with the dataset identifier PXD081718 ^61^. The deep learning segmentation models developed in this study are publicly available through the arivis Cloud platform. Model #1, which performs PT and glomerular segmentation and classification from megalin immunohistochemistry images, is available under access token ME_tlFp2hb4TbXz76qwoZxF457XpmShou724aKOssKU. Model# 2, which was custom trained for semantic segmentation of mouse kidney immunofluorescence images stained for megalin, Na,K-ATPase, and DAPI, is available under access token aoStEkZix4m7hsaOjHkbMYijL29ZvPuLnkZ0G_XkmZA. The dataset will be publicly available upon publication. The imaging datasets generated in this study have been deposited in the BioImage Archive under accession number S-BIAD3843 ^62^. All other data supporting the findings of this study are available from the corresponding author upon reasonable request.

## Acknowledgement

We thank Inger Merete S. Paulsen, Cassandra Bennetzen, Tina Drejer, Christian V. Westberg, and Anna Louise Venning Grandjean for their technical support. The mass spectrometry equipment utilized in this study is part of the Danish Single-Cell Examination Platform (CellX) established with support from the Danish Research Agency Infrastructure Program (5229-0009B). Funding for this project to R.A. Fenton is provided by the Novo Nordisk Foundation (NNF21OC0067647, NNF24OC0095846 and NNF20SA0061466), the Aarhus University Research Foundation (AUFF-E-2022-9-22), the Danish Council for Independent Research (0134-00018B and 3101-00136B) and the Carlsberg Foundation (CF21-0243 and CF23-1530).

## Author contributions

Conceptualization: R.A.F., X.Z. M.R.; Methodology: X.Z., Q.W., M.A., L.K.R., R.A.F.; Data curation: X.Z., M.A., L.K.R., Q.W.; Formal analysis: X.Z.; Visualization: X.Z.; Project administration: R.A.F., X.Z., M.R.; Funding acquisition: R.A.F., M.R.; Writing original draft: X.Z. All authors reviewed and edited the manuscript.

## Declaration of generative AI and AI-assisted technologies in the writing process

During preparation of this work, the authors used Anthropic Claude to assist with editing the manuscript. After using this tool, the authors reviewed and edited the content as needed and take full responsibility for the content of the publication.

