## Supplementary Information for "Deep visual proteomics reveals distinct proximal tubular and glomerular injury programs in experimental diabetic kidney disease"

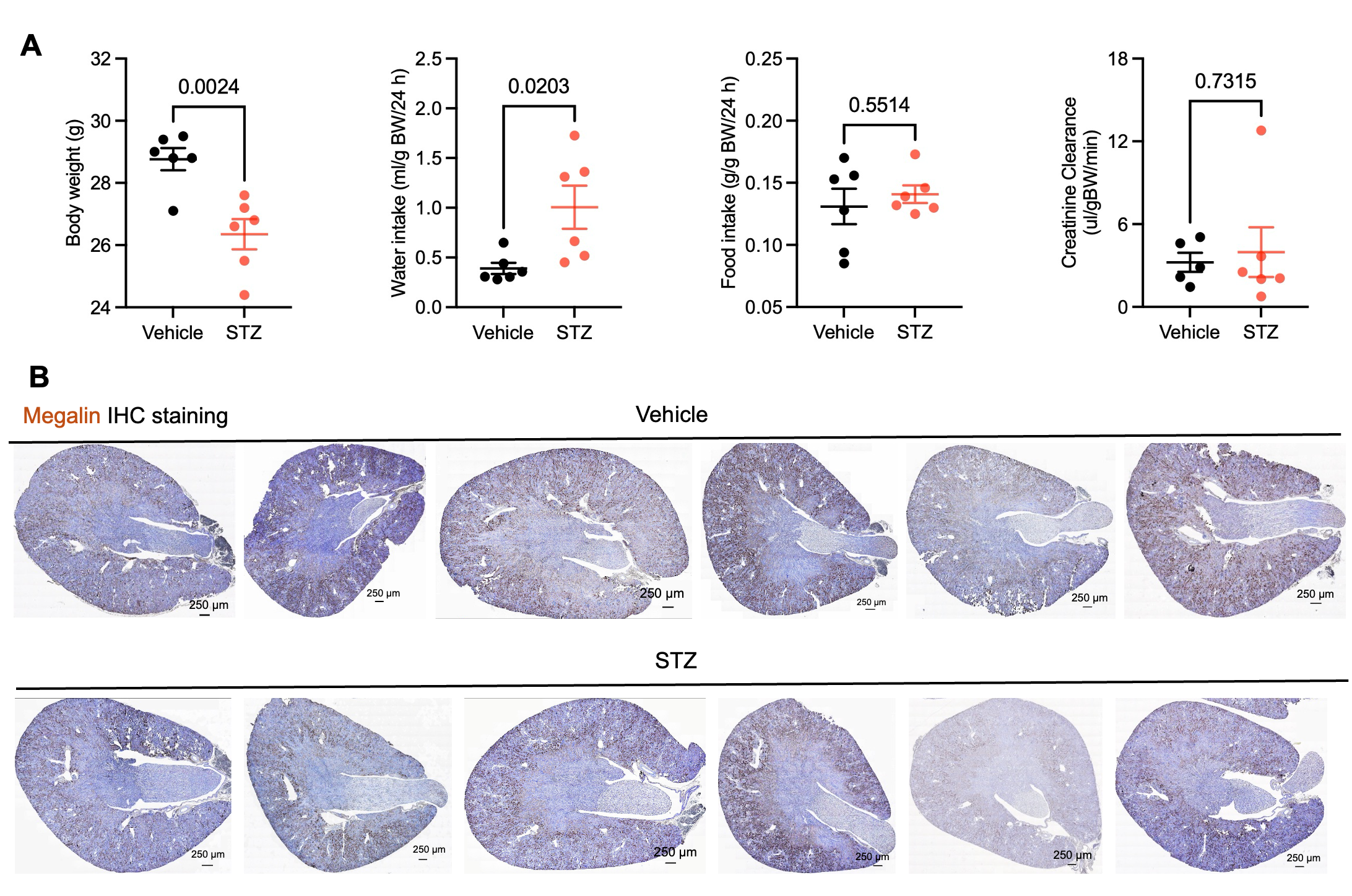

**Supplementary Figure 1. Physiological characterization of the STZ model and anatomical basis for proximal tubule microdissection.** (A) Body weight, water intake, food intake and creatinine clearance. Bars show mean ± SEM; statistical comparisons by unpaired *t test*. (B) Representative megalin immunohistochemistry (IHC) delineating the proximal tubule compartment in kidney sections, used as the anatomical basis for compartment-specific microdissection (Figure 2).

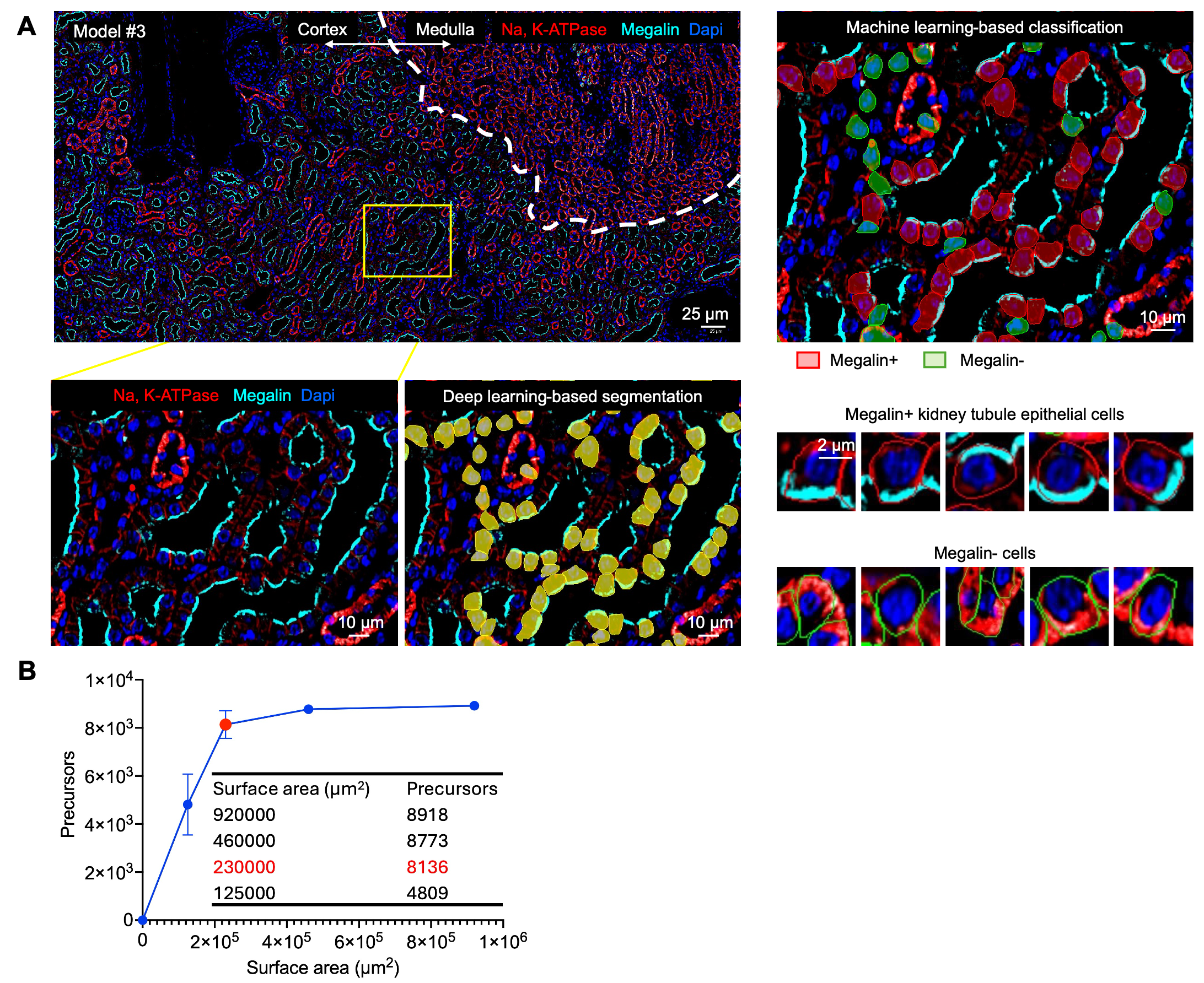

**Supplementary Figure 2. Extending the DVP pipeline to single-cell resolution and defining minimum sample input for proteomic identification.** (A) A cytoplasmic single cell segmentation model combined with a machine learning classifier identifies individual PT epithelial cells, extending the pipeline to single cell resolution. (B) Mass spectrometry precursor (parent ion) identifications as a function of microdissected surface area. Identification rates plateaued above 2.3 × 10^5^ µm², the minimum surface area applied for sample acquisition throughout this study.

| **Supplementary Table 1.** Sensitivity of marker prioritization score to alternative weighting schemes | | | |
| --- | --- | --- | --- |
| Metric | Median | Min | Max |
| Top-20 overlap with base ranking | 18/20 | 3/20 | 20/20 |
| Jaccard similarity (top 20) | 0.818 | 0.081 | 1.000 |
| Spearman rank correlation (full ranking) | 0.989 | 0.956 | 1.000 |
| Sensitivity of composite marker prioritization score to 220 alternative weighting schemes (see Supplementary Methods for details). Overlap and Jaccard similarity are relative to the top 20 candidates under the base weighting (0.35/0.30/0.20/0.15); Spearman rho reflects rank correlation across the full candidate list. | | | |
